## Supplemental information for "Fundamentals of Vaping-Associated Pulmonary Injury Leading to Severe Respiratory Distress"

Supplemental Methods, Figure Legends, and Tables

#### Methods

Histological staining. Lungs of e-cigarette-exposed and control mice were harvested by inflation with formalin. Right lung lobes were sutured off at the right primary bronchus and frozen back in liquid nitrogen immediately following dissection, while the left lobe was manually inflated over the course of 15 minutes with approximately 1.5 mL of formalin until visual confirmation of sufficient inflation. The left lobe was then dissected and submerged in formalin for 24 hours followed by tissue processing, paraffin embedding, and sectioning. The lungs were compressed to attain the primary bronchial tree in the same plane of view as parenchyma. Sections were stained with Harris Hematoxylin and Eosin-Phloxine (H&E) in addition to Movat's Pentachrome reagents to visualize morphometric and structural changes. Lung sections were stained following the Modified Russell-Movat Pentachrome Stain protocol to visualize changes in collagen, elastic fibers, and mucin deposition. Pentachrome staining is interpreted as elastic fibers (black to blue/black), nuclei (blue/black), collagen (yellow to red), reticular fibers (yellow), mucin (bright blue), fibrin (bright red), muscle (red). Trichrome Stain (Masson) Kit (Sigma-Aldrich, catalog #HT15) was used to visualize collagen deposition in hearts and lungs according to manufacturer's protocol. Images were acquired using a Leica DMIL6000 microscope running XY stage tile scanning and subsequently stitched using ImageJ software. Periodic Acid Schiff (PAS) Stain: Slides were deparaffinized and rehydrated following generic procedures. PAS stain was performed following kit protocol specifications (Sigma-Aldrich 1.01646.0001). Hematoxylin solution modified according to Gill III was substituted, as recommended by the protocol, for Hematoxylin solution modified according to Gill II (Sigma-Aldrich GHS216). Finally, slides were mounted with Toluene Solution (Fisher Scientific Lot#103929). Images were acquired using a Leica DMIL6000 microscope. All images were taken in the upper airway as close as possible to the branchpoint of the primary and secondary bronchi.

Immunohistochemistry and confocal microscopy. Paraffin sections (5µm) were deparaffinized, subjected to antigen retrieval in 10mM citrate pH 6.0 and quenched for endogenous autofluorescent activity in 3% Sodium Borohydride (Aldrich 452882-250) in TN buffer (150mM NaCl, 100mM Tris pH 7.6) for 30 minutes. Blocking was performed for 30 minutes at room temperature with 10% Horse Serum in 1xTN. Tissues were immunolabeled with primary antibodies as listed in Supplemental Table 1 overnight at 4°C. Fluorescently conjugated secondary antibodies were diluted 1:200 (Donkey anti-goat 488 Life Tech cat# A11055, Donkey anti-rabbit 555 Invitrogen cat# A32794, Donkey anti-rat 555 Jackson Labs cat# 712-166-153, Donkey anti-chicken 555 Jackson Labs cat# 703-165-155, Donkey anti-rabbit 647 Invitrogen cat# A31573, Donkey anti-chicken 647 Jackson Labs cat# 703-505-155). DAPI (Thermo Fisher cat# 62248) was applied in the final wash step at 0.1µg/ml to label nuclei. Images were acquired using a Leica SP8 confocal microscope and processed with Leica and Photoshop software. Stains were performed on at least three different samples per exposure group, and one technical replicate. Tissue was imaged from trachea, conducting airways, and lung parenchyma. Muc5ac Immunostain: Frozen sections prepared identical to those used for spatial transcriptomics (see below) were used to visualize Muc5ac. Slides stored at -80°C were warmed to room temperature and washed twice in PBS (.37 M NaCl, 27 mM KCl, 100 mM $Na_2HPO_4$, 18 mM $KH_2PO_4$), then fixed in -20°C methanol for 30 minutes. Two more washes in PBS were performed, then tissue was blocked using 10% Horse Serum for 30 minutes. Next, two washes in PBS were followed by immunolabeling with Muc5ac (U.S. Biotech cat# 1364248) and ECAD (R&D cat# AF748) overnight at 4$^{o}$C. Two more washes in PBS were done the following day, and fluorescently conjugated secondary antibodies were used to detect primary antibodies (Donkey anti-Rabbit 555 Jackson Labs 703-165-155, Donkey anti-Goat Life Tech A11055) for one hour and thirty minutes at room temperature. After two washes in PBS, DAPI (Thermo Fisher cat# 62248) was applied in the final wash step at 0.1µg/ml to label nuclei for 15 minutes in PBS, and slides were coverslipped and mounted in Vectashield (Vector labs cat# H-1000). Images were acquired using LeicaSP8 confocal microscope and images were processed with Leica software. Tissue was imaged in most distal airways where mucin secretion is uncommon.

<u>Immunoblotting</u>. Lungs from vaped animals and air-exposed controls were collected and stored at -80 °C for future analysis. Frozen lung tissues were homogenized in lysis buffer containing protease and phosphatase inhibitor cocktails (Sigma-Aldrich; P8340, P5726, P2850). Bradford assay was performed to analyze and normalize protein concentrations and lysates were prepared by addition of NuPAGE LDS Sample Buffer (ThermoFisher; NP0007) and 100 $\mu$M dithiothreitol (Bio-Rad; 161-0611). Samples were sonicated and boiled for 5 minutes at 95 °C and stored at -80 °C. Proteins were separated on a 4–12% NuPAGE Bis-Tris gel (ThermoFisher; NP0321BOX) and transferred onto a polyvinylidene fluoride membrane. Nonspecific binding sites were blocked using Odyssey blocking buffer (LICOR, 927-60001) and proteins were labeled with primary antibodies in 0.2% Tween in Odyssey blocking buffer overnight. After multiple washes, blots were incubated with secondary antibodies in 0.2% Tween 20 in Odyssey blocking buffer for 1.5 hours at room temperature and scanned using the LICOR Odyssey CLx scanner. Quantification was performed using ImageJ software. Antibodies and their dilutions are listed in Supplemental Table 2.

<u>Spatial transcriptomic analyses</u>. <u>*Sample preparation*</u>. Lungs were inflated through the trachea with a 1:1 PBS/OCT mix (1X PBS, Corning Cat# 21-040-CV; TissueTek O.C.T. Compound, VWR Cat# 25608-930) with RNAse inhibitor at 0.2 U/$\mu$l (Millipore Sigma, Cat# 3335399001). Tissues were embedded in OCT and flash frozen using an isopentane and liquid nitrogen bath. Tissue sections 10 $\mu$m were obtained on a CryoStar NX70 (ThermoScientific) and processed immediately for spatial transcriptional analysis or stored for histological stains. Blocks and sections were maintained at -80C for long-term storage. <u>*Image collection and spatial transcriptomic library preparation.*</u> Freshly obtained cryosections were placed in Visium gene expression slides (10X Genomics, Cat# 2000233) for processing. Tissue staining with hematoxylin and eosin and image collections were performed as recommended by the Visium protocol. Images were collected on a Leica DMI6000 B on a 5X objective at a 1.16 $\mu$m/pixel capture resolution (Supplemental Figure 2). Spatial transcriptomic libraries were prepared using Visium Spatial Gene Expression Slide & Reagent Kit following manufacturer's protocol (10X Genomics, PN-1000184). Lung permeabilization time was optimized Visium Spatial Tissue Optimization Slide & Reagent Kit (10X Genomics, PN-1000193). Samples were processed together to avoid introduction of technical batch effects. Library concentration and fragment size distribution of each library were tested with Bioanalyzer (Agilent High Sensitivity DNA Kit, Cat. # 5067-4626; average library size: 500-610 bp). The sequencing libraries were quantified by quantitative PCR (KAPA Biosystems Library Quantification Kit for Illumina platforms P/N KK4824) and Qubit 3.0 with dsDNA HS Assay Kit (Thermo Fisher Scientific). Sequencing libraries were submitted to the UCSD IGM Genomics Core for sequencing (NovaSeq 6000), aiming for > 50K reads per spot.

<u>Cardiomyocyte Cross Sectional Area Quantitation</u>: Heart sections of all treatment groups were acquired and stained as previously described with the exception that heart samples were not

treated with sodium borohydride. 24 images of each right ventricle were taken using a Leica SP8 confocal microscope at a 400x magnification. Quantification of the cross sectional area of right ventricular myocytes was done in ImageJ software using a pixel/um ratio of 3.5 for all images analyzed. Myocytes that contained two clear intercalated discs at each end with an associated nucleus were measured by drawing a line from one intercalated disc to the other, then measuring the length of the line. All measurements taken from each experimental group are listed in the table below. Data points were input into Prism5 software to compose a graph and run a one way anova test using Kruskal-Wallis metrics, $p<0.01$. Male Vape (n=4, 133 measurements, SD: +/-12.9 um). Female Vape (n=4, 116 measurements, SD: +/-14.13). Male No Vape (n=3, 83 measurements, SD: +/-14.97). Female No Vape (n=3, 72 measurements, SD: +/-9.935).

Statistics. For mean linear intercept and bronchiole wall area percentage, unpaired t-tests were performed between the vape and no vape groups with a 95% confidence interval and a two-tailed p value of 0.0167 for 1D and 0.0098 for 1E I using GraphPad Prism Version 5.02. For echocardigraphy, 5-7 mice per group ANOVA with Kruskal-Wallis significant differences test with $p<0.05$(*), $p<0.01$(**), $p<0.001$(***). For differential expression analysis, Wilcoxon rank sum test was performed with selection for a threshold of 0.05 for an adjusted p-value and a log (FC) >0.25 was used to define statistically significant and differentially expressed genes (DEGs).G0 term analysis was performed with p-value cutoff of 0.05 using Benjamini-Hochberg Procedure. For immunoblot analysis, two-tailed unpaired t-test was used to compare 2 groups of vape and non-vape samples. Statistical analysis was performed using GraphPad Prism. A *p*-value of $<0.05$ was considered statistically significant. For cardiomyocyte cross-sectional length, right ventricular myocytes were analyzed in male and female, vaped and non-vaped samples. A one way anova using Kruskal-Wallis t-test metrics was performed on the total number of measurements per group ($p<0.01$). Male Vape (n=4, 133 measurements, SD: +/-12.9 um). Female Vape (n=4, 116 measurements, SD: +/-14.13). Male No Vape (n=3, 83 measurements, SD: +/-14.97). Female No Vape (n=3, 72 measurements, SD: +/-9.935).

| Marker | Species | Company | Catalog # | Dilution |
|---|---|---|---|---|
| ECAD | Goat | R&D Systems | AF748 | 1:100 |
| Vimentin | Chicken | Invitrogen | PA1-10003 | 1:400 |
| CD11C | Rabbit | Cell Signaling Tech | 97585S | 1:200 |
| RAGE | Goat | R&D Systems | AF1145 | 1:100 |
| KRT5 | Chicken | Biolegend | 905901 | 1:200 |
| AQ5 | Rabbit | Abcam | Ab78486 | 1:400 |
| CD9 | Rabbit | Abcam | Ab92726 | 1:100 |
| Alpha Tubulin | Rat | Biorad | 1703 | 1:400 |
| Muc5ac | Rabbit | US Biotech | 1364248 | 1:100 |
| Desmin | Rabbit | Abcam | Ab15200 | 1:200 |

Supplemental Table 1. Antibodies used for immunohistochemistry.

| Marker | Species | Company | Catalog # | Dilution |
|---|---|---|---|---|
| CD11b | Rabbit | Abcam | Ab133357 | 1:500 |
| CD11c | Rabbit | Cell Signaling | 97585s | 1:500 |
| CD206 | Rabbit | Abcam | Ab64693 | 1:500 |
| CD45 | Goat | R&D | AF114 | 1:500 |
| E-Cadherin | Rabbit | R&D | AF748 | 1:500 |
| Fibronectin | Rabbit | Sigma | F3648 | 1:250 |
| GAPDH | Goat | Sicgen | AB0067 | 1:3000 |
| HMGB1 | Rabbit | Abcam | Ab18256 | 1:250 |
| IL1-β | Rabbit | Invitrogen | P420B | 1:50 |
| IL-6 | Mouse | Sino Biological | 10395-MM19 | 1:250 |
| MUC1 | Rabbit | Abcam | Ab109185 | 1:250 |
| MUC5AC | Rabbit | My Biosource | MBS2028179 | 1:200 |

Supplemental Table 2. Antibodies used for immunoblotting.

| | No Vape Female | No Vape Male | Vape Female | Vape Male |
|---|---|---|---|---|
| EF | 74.82± 0.464655 | 74.99±2.531607 | 64.95±3.74678 | 59.43±2.98888 |
| FS | 42.24±0.476157 | 42.66±2.090859 | 34.4±2.743135 | 30.66±1.985322 |
| LV_Mass | 81.53±18.11191 | 89.68±13.87718 | 80.95±4.805302 | 88.3±11.93415 |
| LV_Vold | 35.42±6.053918 | 42.03±7.525448 | 36.39±4.106111 | 44.07±6.537644 |
| LV_Vols | 8.916±1.502419 | 10.612±2.718028 | 12.75±1.988373 | 17.92±3.238416 |
| LVIDd | 3.007±0.212416 | 3.225±0.239866 | 3.045±0.139681 | 3.291±0.207086 |
| LVIDs | 1.736±0.115514 | 1.852±0.185913 | 1.997±0.119995 | 2.283±0.167831 |
| IVSd | 0.9405±0.08526 | 0.957±0.042974 | 0.8723±0.048719 | 0.9123±0.022281 |
| IVSs | 1.355±0.098609 | 1.382±0.093474 | 1.3±0.060127 | 1.265±0.108597 |
| LVPWd | 0.75±0.171771 | 0.726±0.035867 | 0.8028±0.043058 | 0.718±0.111153 |
| LVPWs | 0.8085±0.184679 | 0.7938±0.072507 | 0.8578±0.063111 | 0.7594±0.064659 |

Supplemental Table 3. Echocardiographic measurements shown by experimental group and gender. Values represent mean ± standard deviation.

| Male Vape | | Female Vape | | Male No Vape | | Female No Vape | |
|---|---|---|---|---|---|---|---|
| Sample # | Measurements | Sample # | Measurements | Sample # | Measurements | Sample # | Measurements |
| 21053 | 25 | 21075 | 23 | 21061 | 29 | 21071 | 24 |
| 21099 | 38 | 21077 | 28 | 21042 | 19 | 21041 | 20 |
| 21044 | 43 | 21043 | 34 | 21113 | 35 | 21059 | 28 |
| 21065 | 27 | 21089 | 31 | | | | |
| Total Measurements: 133 | | Total Measurements: 116 | | Total Measurements: 83 | | Total Measurements: 72 | |
| Mean: 62.51 | | Mean: 60.93 | | Mean: 56.92 | | Mean: 53.76 | |
| % Increase: 8.94 | | % Increase: 11.77 | | | | | |

**Supplemental Table 4. Parameters for cardiomyocyte length measurements.**
**Supplemental Figures and Legends**.

### Supplemental Figure 1

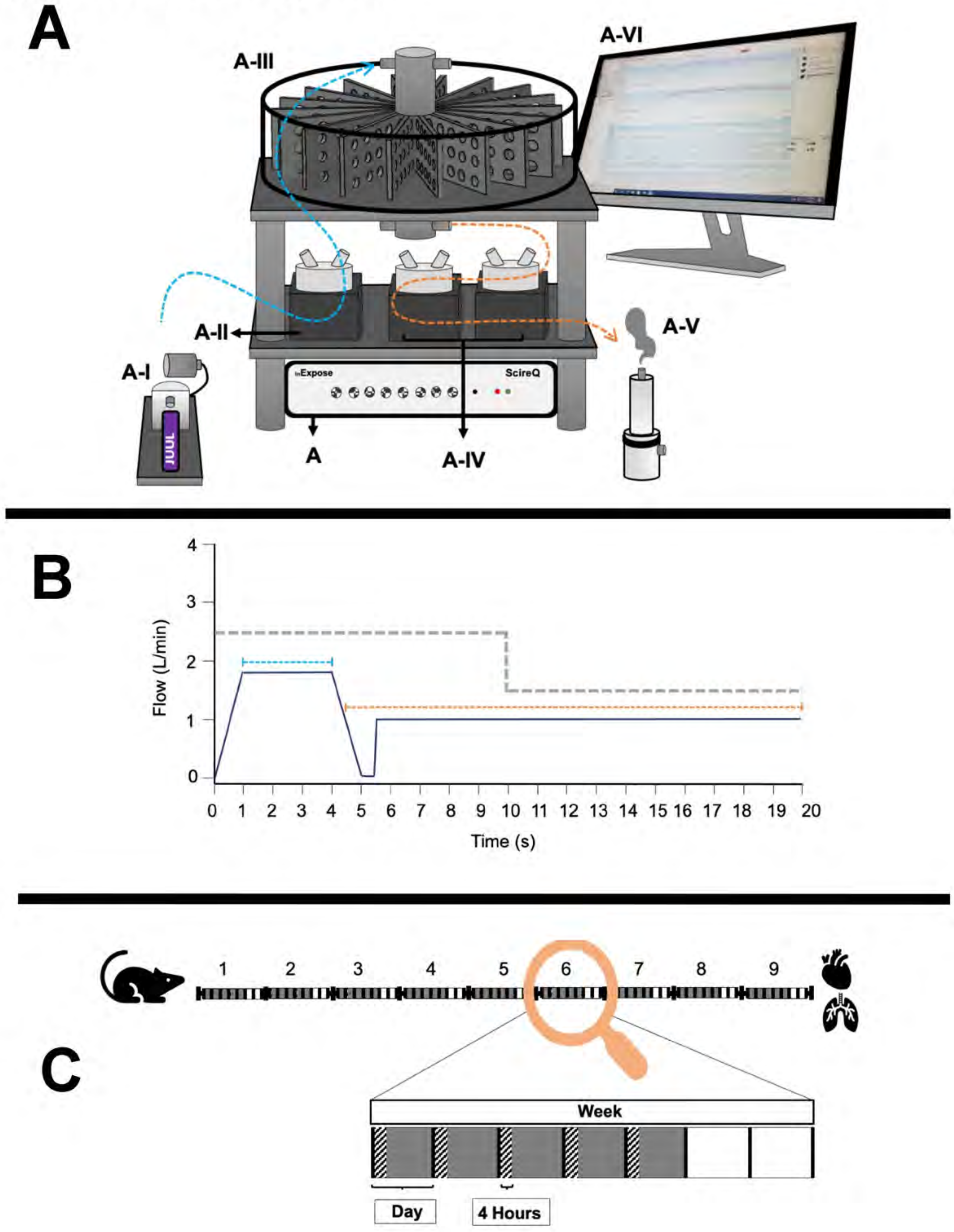

Supplemental Figure 1. Vaping protocol details showing InExpose equipment (**A**), vaping topography profile (**B**), and time course of exposure schematic (**C**). InExpose base unit is software controlled to regulate exposure time and duration (A). ENDS device stroke controller ensures consistent e-vapor deposition (A-I). InExpose pump 1 generates e-vapor puffs (A-II; one pump per exposure chamber, only one pump shown). Whole body chamber houses up to 16 subjects for simultaneous exposure (A-III). InExpose exhaust pumps 2 and 3 (A-IV; one pump per exposure chamber). Buffer chamber pulls vapor from exposure chamber in order to circulate fresh air during inter-puff intervals (A-V). FlexiWare software controls the base unit to regulate vaping topography time and duration (A-VI). (**B**) Schematic of vaping topography. Mice in exposure chambers received e-vapor delivered in 3 second puffs (blue dashed line; 1.8 L/min flow rate) in 20 second intervals (orange dashed line; 1 L/min air flow rate). Exhaust pump flow rate alternated between puff and interval at rates of 2.5 L/min and 1.5 L/min, respectively (grey dashed line). (**C**) C57BL/6J mice were exposed to Organx Peach Ice flavored vape juice delivered as e-vapor from JUUL pens. Chamber habitation for 4 hours per day, 5 days per week over 9 week time course. Vaping Days (gray boxes), Exposure time (hatched rectangles), and Rest Days (white boxes) are shown.

### Supplemental Figure 2

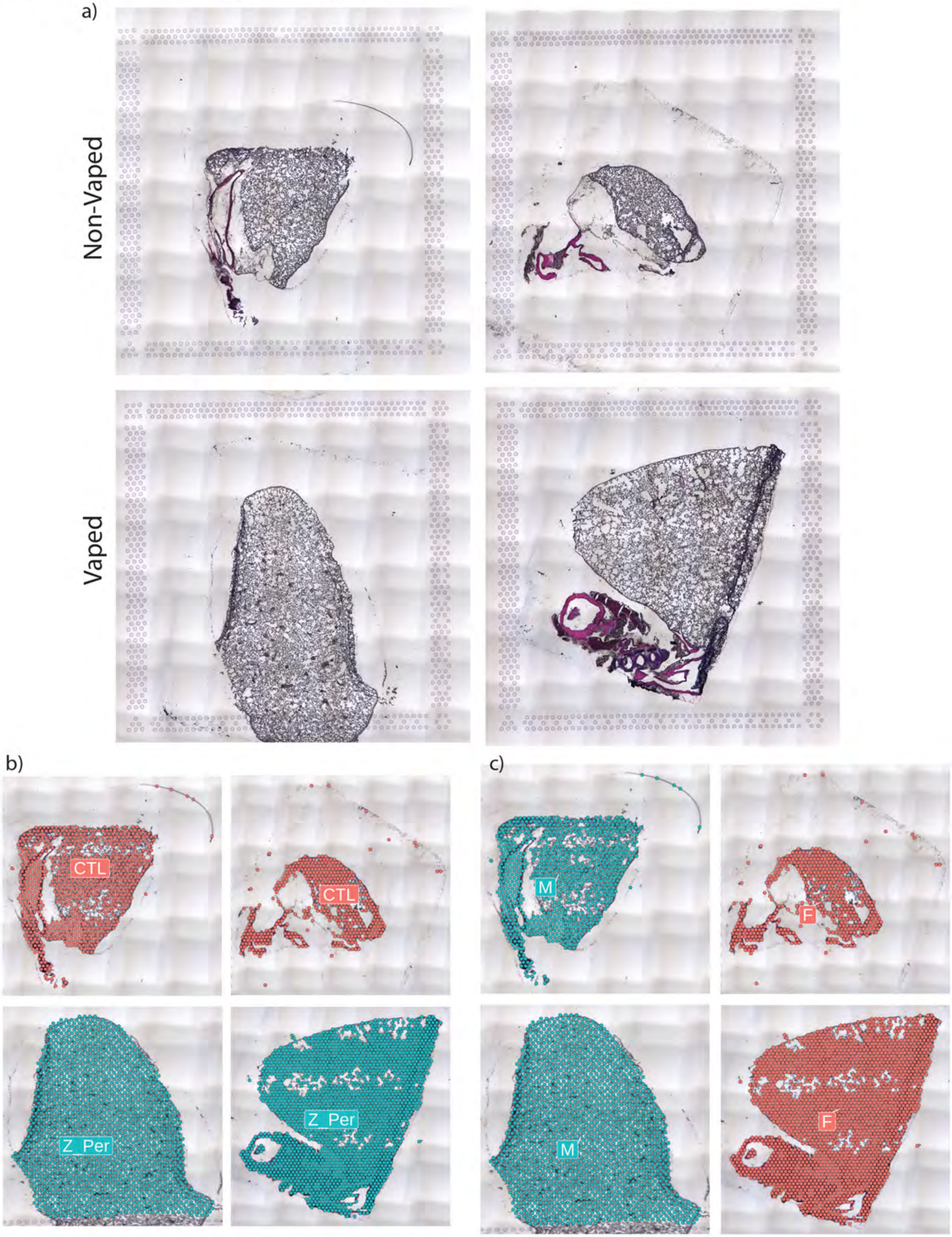

Supplemental Figure 2. Input for Visium Spatial transcriptomic experiment. a) Hematoxylin and Eosin (H&E) micrographs of Non-Vaped and Vaped samples. Identification of transcriptional spots overlaid on H&E micrographs of Non-Vaped and Vaped samples, classified by b) treatment and c) sex of collected sample.

### Supplemental Figure 3

10x GENOMICS Space Ranger • count

#### CTL_1_M63

Summary | Analysis

**842**
Number of Spots Under Tissue

**211,852**
Mean Reads per Spot

**3,792**
Median Genes per Spot

##### Sequencing

| | |
|---|---|
| Number of Reads | 178,379,315 |
| Valid Barcodes | 96.2% |
| Valid UMIs | 99.6% |
| Sequencing Saturation | 89.3% |
| Q30 Bases in Barcode | 93.9% |
| Q30 Bases in RNA Read | 88.8% |
| Q30 Bases in UMI | 93.7% |

##### Mapping

| | |
|---|---|
| Reads Mapped to Genome | 86.2% |
| Reads Mapped Confidently to Genome | 82.5% |
| Reads Mapped Confidently to Intergenic Regions | 7.2% |
| Reads Mapped Confidently to Intronic Regions | 3.0% |
| Reads Mapped Confidently to Exonic Regions | 72.3% |
| Reads Mapped Confidently to Transcriptome | 70.8% |
| Reads Mapped Antisense to Gene | 0.5% |

##### Spots

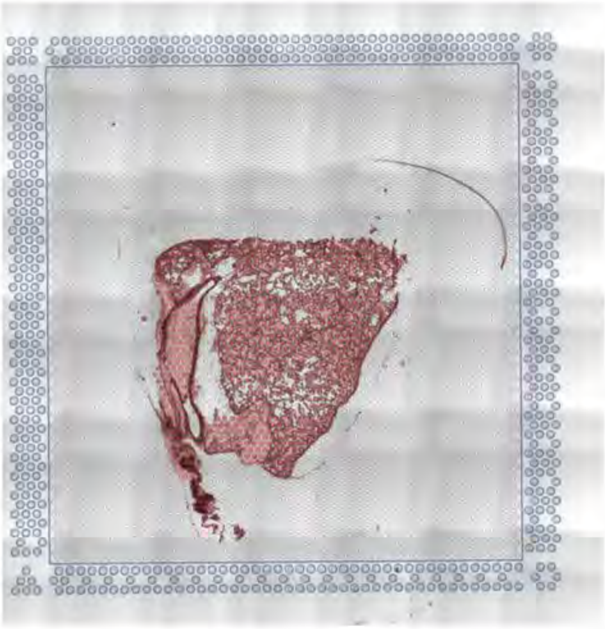

| | |
|---|---|
| Fraction Reads in Spots Under Tissue | 76.8% |
| Mean Reads per Spot | 211,852 |
| Mean Reads Under Tissue per Spot | 150,745 |
| Median Genes per Spot | 3,792 |
| Total Genes Detected | 18,057 |
| Median UMI Counts per Spot | 9,741 |

##### Sample

| | |
|---|---|
| Sample ID | CTL_1_M63 |
| Sample Description | |
| Chemistry | Spatial 3' v1 |
| Slide Serial Number | V19N11-097-A1 |
| Reference Path | …refdata-gex-mm10-2020-A |
| Transcriptome | mm10-2020-A |
| Pipeline Version | spaceranger-1.2.2 |

Supplemental Figure 3. Alignment quality control for Visium Spatial transcriptomic experiment.
SpaceRanger 1.2.2 quality control summary for Non-Vaped Male.

### Supplemental Figure 4

10x GENOMICS Space Ranger • count

#### CTL_2_F62

Summary Analysis

**473**
Number of Spots Under Tissue

**370,210**
Mean Reads per Spot

**3,979**
Median Genes per Spot

##### Sequencing

| | |
|---|---|
| Number of Reads | 175,109,131 |
| Valid Barcodes | 96.7% |
| Valid UMIs | 99.8% |
| Sequencing Saturation | 92.1% |
| Q30 Bases in Barcode | 94.1% |
| Q30 Bases in RNA Read | 88.3% |
| Q30 Bases in UMI | 93.9% |

##### Mapping

| | |
|---|---|
| Reads Mapped to Genome | 91.4% |
| Reads Mapped Confidently to Genome | 87.1% |
| Reads Mapped Confidently to Intergenic Regions | 5.1% |
| Reads Mapped Confidently to Intronic Regions | 2.6% |
| Reads Mapped Confidently to Exonic Regions | 79.4% |
| Reads Mapped Confidently to Transcriptome | 77.8% |
| Reads Mapped Antisense to Gene | 0.5% |

##### Spots

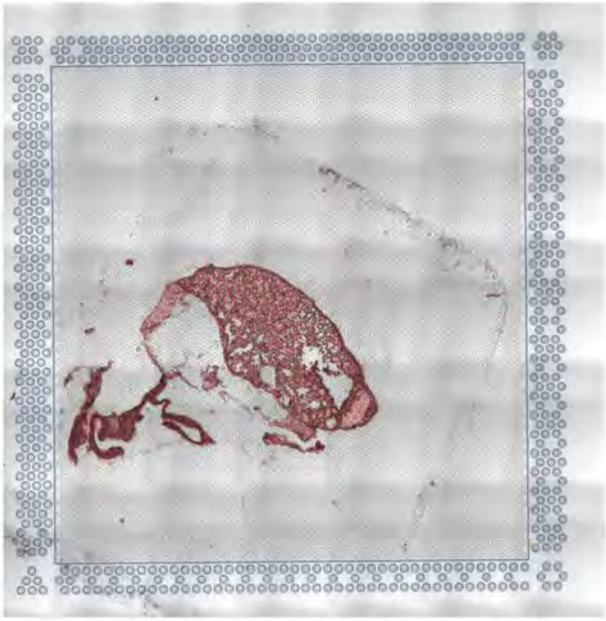

| | |
|---|---|
| Fraction Reads in Spots Under Tissue | 73.6% |
| Mean Reads per Spot | 370,210 |
| Mean Reads Under Tissue per Spot | 252,799 |
| Median Genes per Spot | 3,979 |
| Total Genes Detected | 17,863 |
| Median UMI Counts per Spot | 10,471 |

##### Sample

| | |
|---|---|
| Sample ID | CTL_2_F62 |
| Sample Description | |
| Chemistry | Spatial 3' v1 |
| Slide Serial Number | V19N11-097-B1 |
| Reference Path | …refdata-gex-mm10-2020-A |
| Transcriptome | mm10-2020-A |
| Pipeline Version | spaceranger-1.2.2 |

Supplemental Figure 4. Alignment quality control for Visium Spatial transcriptomic experiment. SpaceRanger 1.2.2 quality control summary for Non-Vaped Female.

### Supplemental Figure 5

10x GENOMICS Space Ranger • count

#### 0_Per_1_M24

Summary | Analysis

**1,548**
Number of Spots Under Tissue

**129,766**
Mean Reads per Spot

**2,917**
Median Genes per Spot

##### Sequencing

| | |
|---|---|
| Number of Reads | 200,877,291 |
| Valid Barcodes | 96.1% |
| Valid UMIs | 99.6% |
| Sequencing Saturation | 89.9% |
| Q30 Bases in Barcode | 94.4% |
| Q30 Bases in RNA Read | 89.1% |
| Q30 Bases in UMI | 94.0% |

##### Mapping

| | |
|---|---|
| Reads Mapped to Genome | 85.3% |
| Reads Mapped Confidently to Genome | 80.4% |
| Reads Mapped Confidently to Intergenic Regions | 4.1% |
| Reads Mapped Confidently to Intronic Regions | 2.6% |
| Reads Mapped Confidently to Exonic Regions | 73.7% |
| Reads Mapped Confidently to Transcriptome | 72.2% |
| Reads Mapped Antisense to Gene | 0.5% |

##### Spots

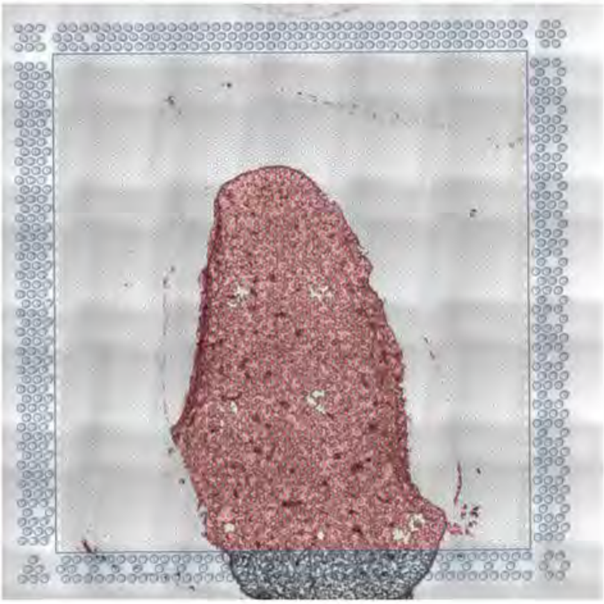

| | |
|---|---|
| Fraction Reads in Spots Under Tissue | 90.6% |
| Mean Reads per Spot | 129,766 |
| Mean Reads Under Tissue per Spot | 111,104 |
| Median Genes per Spot | 2,917 |
| Total Genes Detected | 17,584 |
| Median UMI Counts per Spot | 6,282 |

##### Sample

| | |
|---|---|
| Sample ID | 0_Per_1_M24 |
| Sample Description | |
| Chemistry | Spatial 3' v1 |
| Slide Serial Number | V19N11-097-C1 |
| Reference Path | ...refdata-gex-mm10-2020-A |
| Transcriptome | mm10-2020-A |
| Pipeline Version | spaceranger-1.2.2 |

Supplemental Figure 5. Alignment quality control for Visium Spatial transcriptomic experiment. SpaceRanger 1.2.2 quality control summary for Vaped Male.

### Supplemental Figure 6

10x GENOMICS Space Ranger • count

#### 0_Per_2_F31

Summary Analysis

**2,053**
Number of Spots Under Tissue

**154,228**
Mean Reads per Spot

**4,167**
Median Genes per Spot

##### Sequencing

| | |
|---|---|
| Number of Reads | 316,630,551 |
| Valid Barcodes | 95.2% |
| Valid UMIs | 99.8% |
| Sequencing Saturation | 82.2% |
| Q30 Bases in Barcode | 94.5% |
| Q30 Bases in RNA Read | 90.0% |
| Q30 Bases in UMI | 94.2% |

##### Mapping

| | |
|---|---|
| Reads Mapped to Genome | 88.7% |
| Reads Mapped Confidently to Genome | 85.0% |
| Reads Mapped Confidently to Intergenic Regions | 4.3% |
| Reads Mapped Confidently to Intronic Regions | 2.8% |
| Reads Mapped Confidently to Exonic Regions | 77.9% |
| Reads Mapped Confidently to Transcriptome | 76.0% |
| Reads Mapped Antisense to Gene | 0.7% |

##### Spots

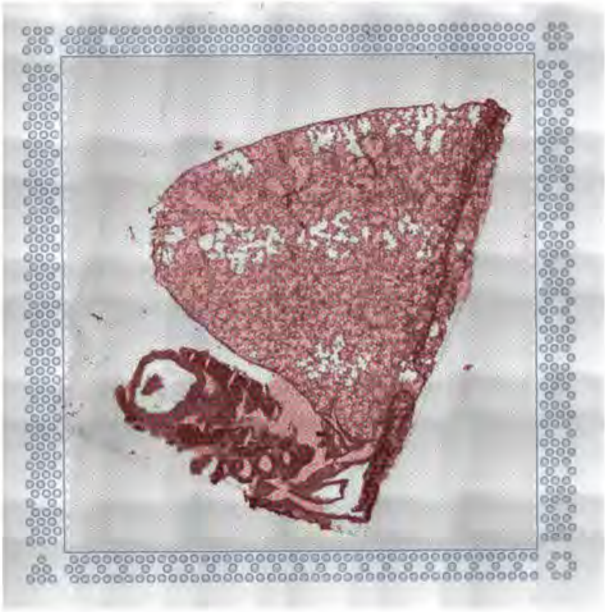

| | |
|---|---|
| Fraction Reads in Spots Under Tissue | 91.7% |
| Mean Reads per Spot | 154,228 |
| Mean Reads Under Tissue per Spot | 133,588 |
| Median Genes per Spot | 4,167 |
| Total Genes Detected | 20,508 |
| Median UMI Counts per Spot | 11,355 |

##### Sample

| | |
|---|---|
| Sample ID | 0_Per_2_F31 |
| Sample Description | |
| Chemistry | Spatial 3' v1 |
| Slide Serial Number | V19N11-097-D1 |
| Reference Path | …refdata-gex-mm10-2020-A |
| Transcriptome | mm10-2020-A |
| Pipeline Version | spaceranger-1.2.2 |

Supplemental Figure 6. Alignment quality control for Visium Spatial transcriptomic experiment. SpaceRanger 1.2.2 quality control summary for Vaped Female.

### Supplemental Figure 7

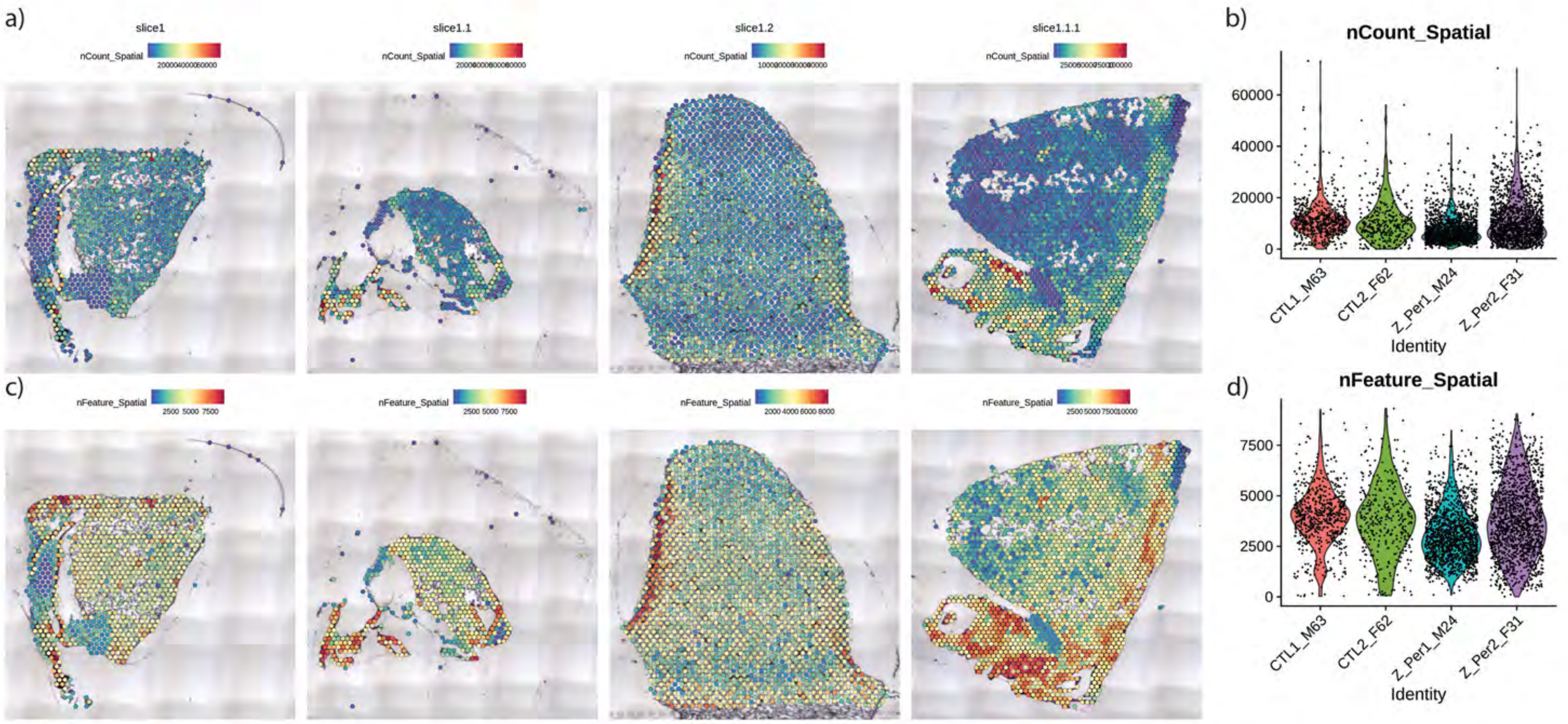

Supplemental Figure 7. Quality control for Visium Spatial transcriptomic experiment. Spatial expression and distribution of a) UMI and c) detected gene counts in samples. Violin plots indicating the spot distribution of b) UMI and d) detected gene counts in samples.

### Supplemental Figure 8

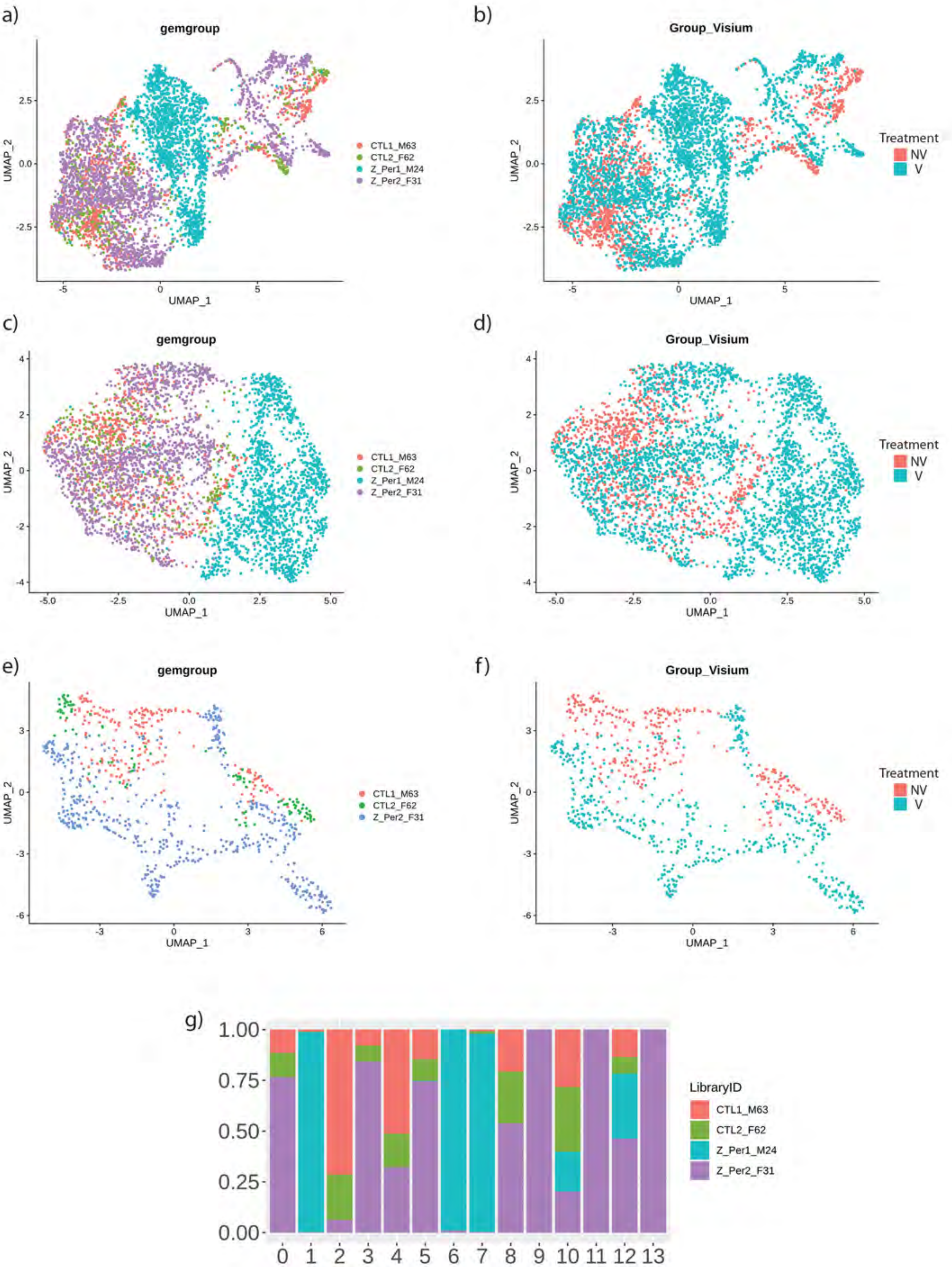

Supplemental Figure 8. Dimensionality reduction reveals transcriptional profiles of Non-Vaped and Vaped pulmonary tissue. Uniform Manifold Approximation and Projection (UMAP) dimensionality reduction projections of spatial transcriptional data color-coded by sample and treatment in global lung tissue (panels a & b), parenchyma (panels c & d) and upper airway (panels e & f). g) Relative spot contributions per sample to each cluster as shown in UMAP (panel a).

### Supplemental Figure 9

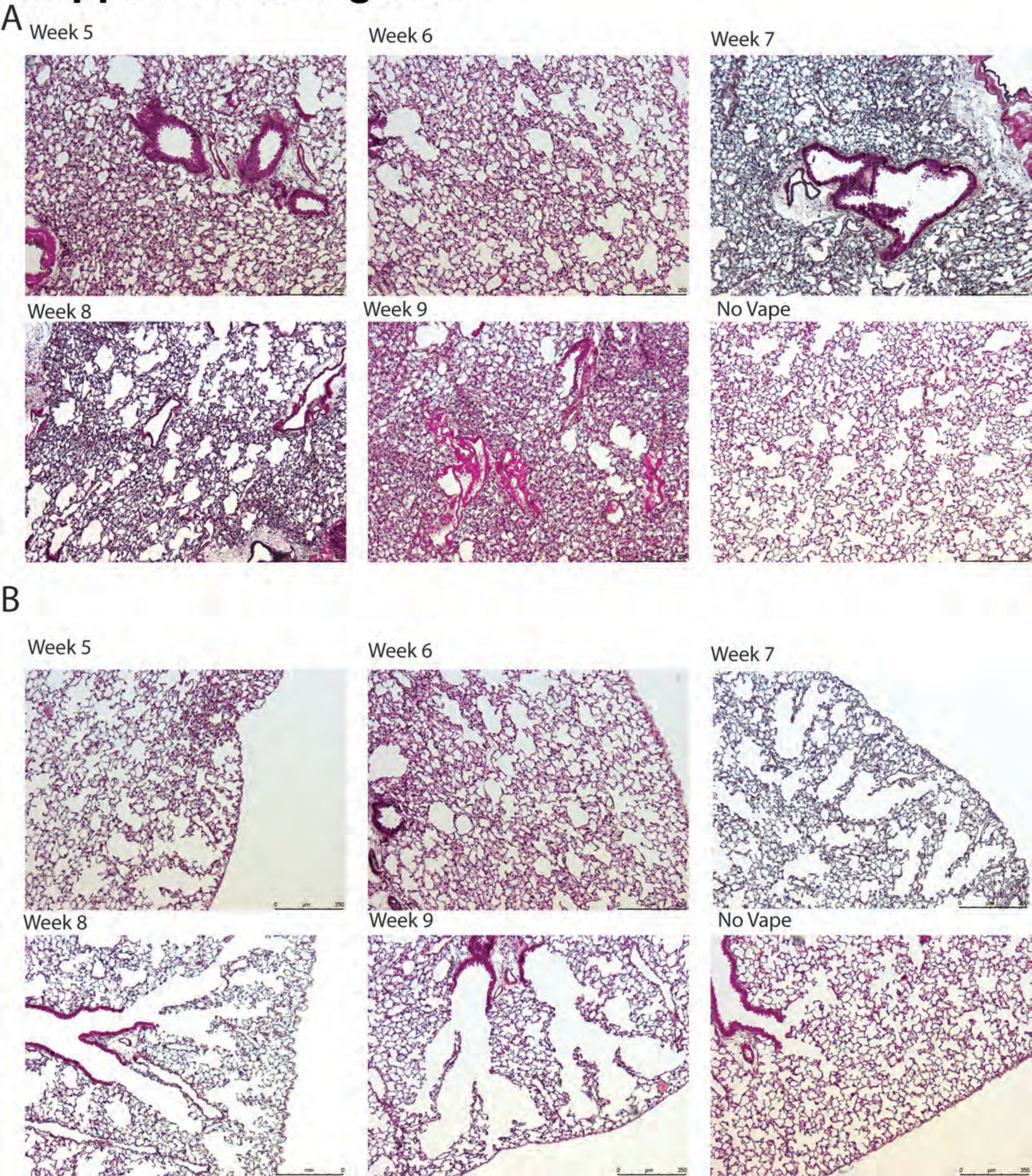

Supplemental Figure 9. Longitudinal time course of vaping injury progression in lungs from Weeks 5 to 9. **(a)** Representative images of lung sections stained with Movat's Pentachrome following 5-9 weeks of vape exposure and no vape for comparison. Time course shows increased cellular density with increased time of vape exposure. **(b)** Representative images of lung sections stained with Masson's Trichrome following 5-9 weeks of vape exposure and no vape for comparison. Time course shows increased alveolar rarefaction with increased time of vape exposure.

### Supplemental Figure 10 (Part 1)

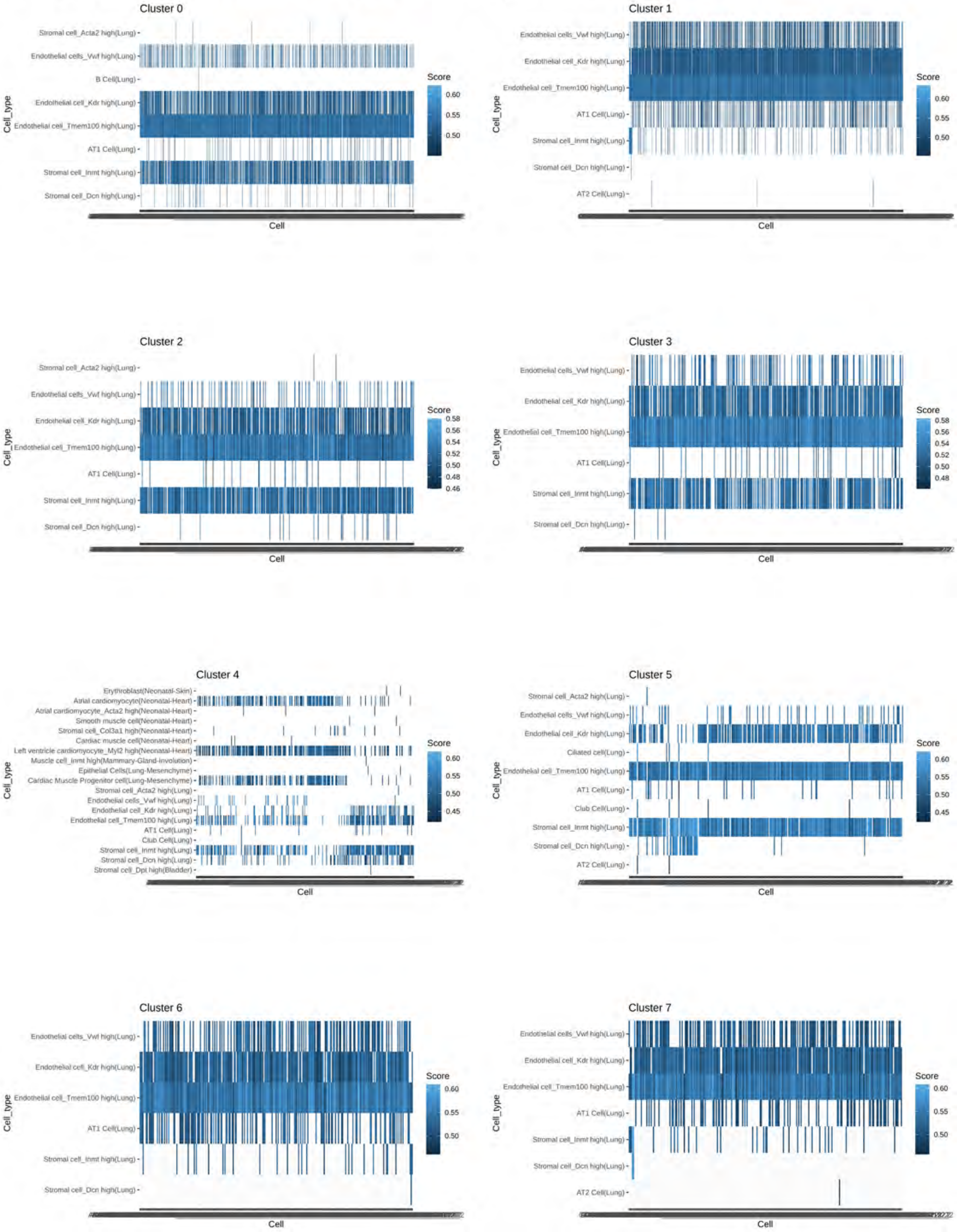

### Supplemental Figure 10 (Part 2)

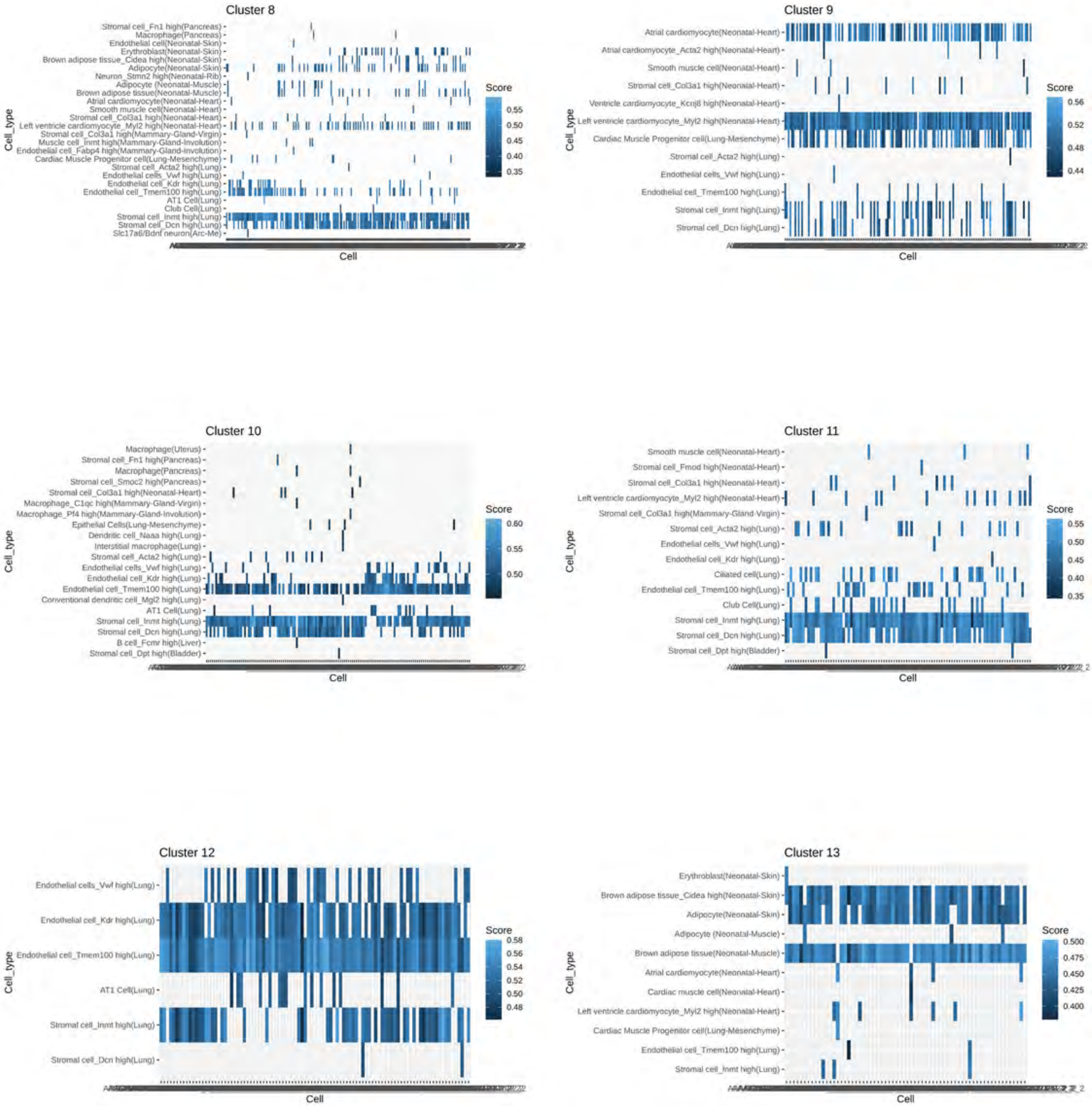

Supplemental Figure 10. Mapping per cluster of spatial transcriptome to MCA database. Heatmap with each row representing mapped cell type and each column a spot within the cluster with mapping score color-coded according to heatmap scale.

### Supplemental Figure 11

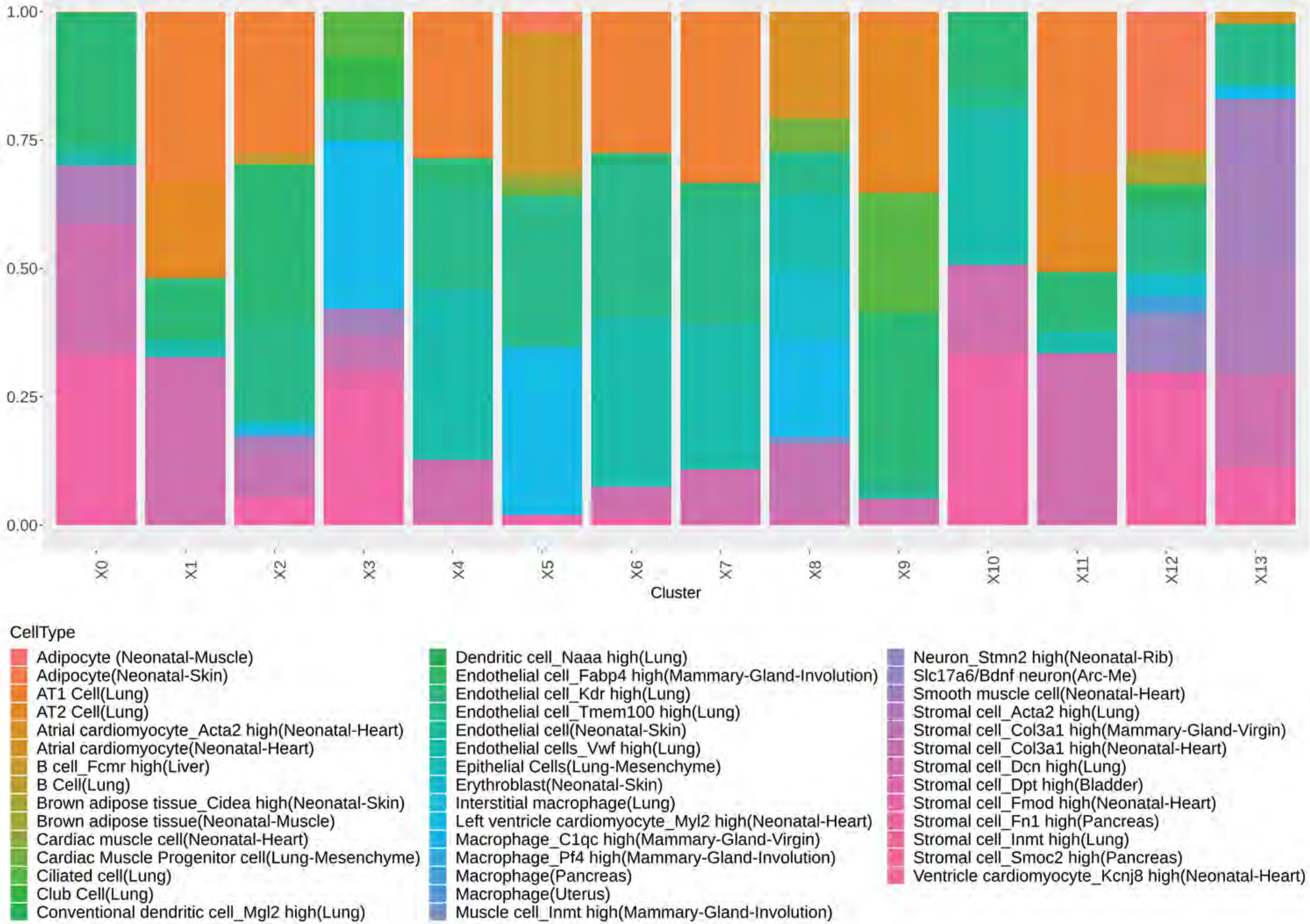

Supplemental Figure 11. Putative cell type contributions by cluster relative to MCA database. (Extended plot).

### Supplemental Figure 12

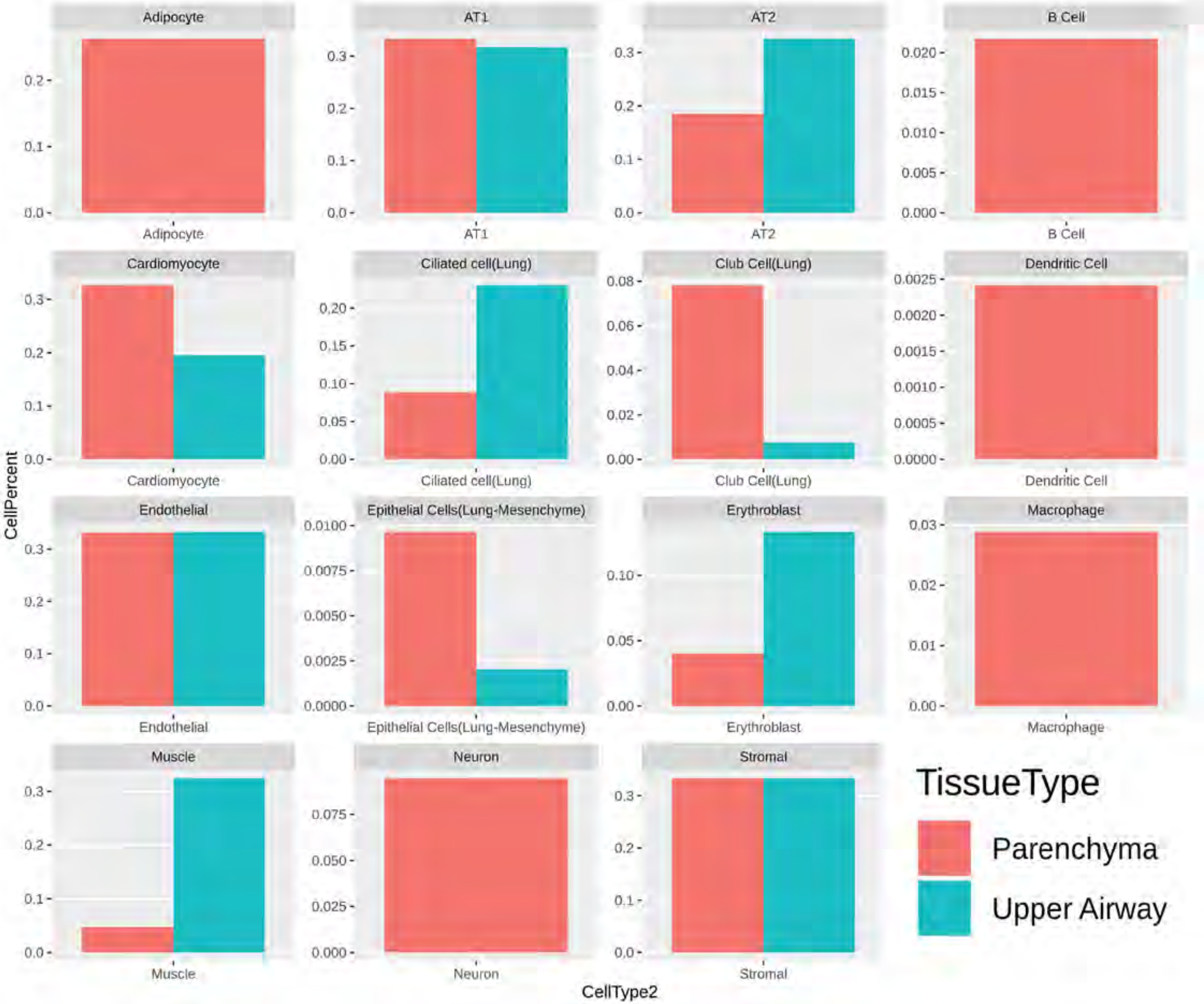

Supplemental Figure 12. Putative cell type contributions by tissue type relative to MCA database.

### Supplemental Figure 13

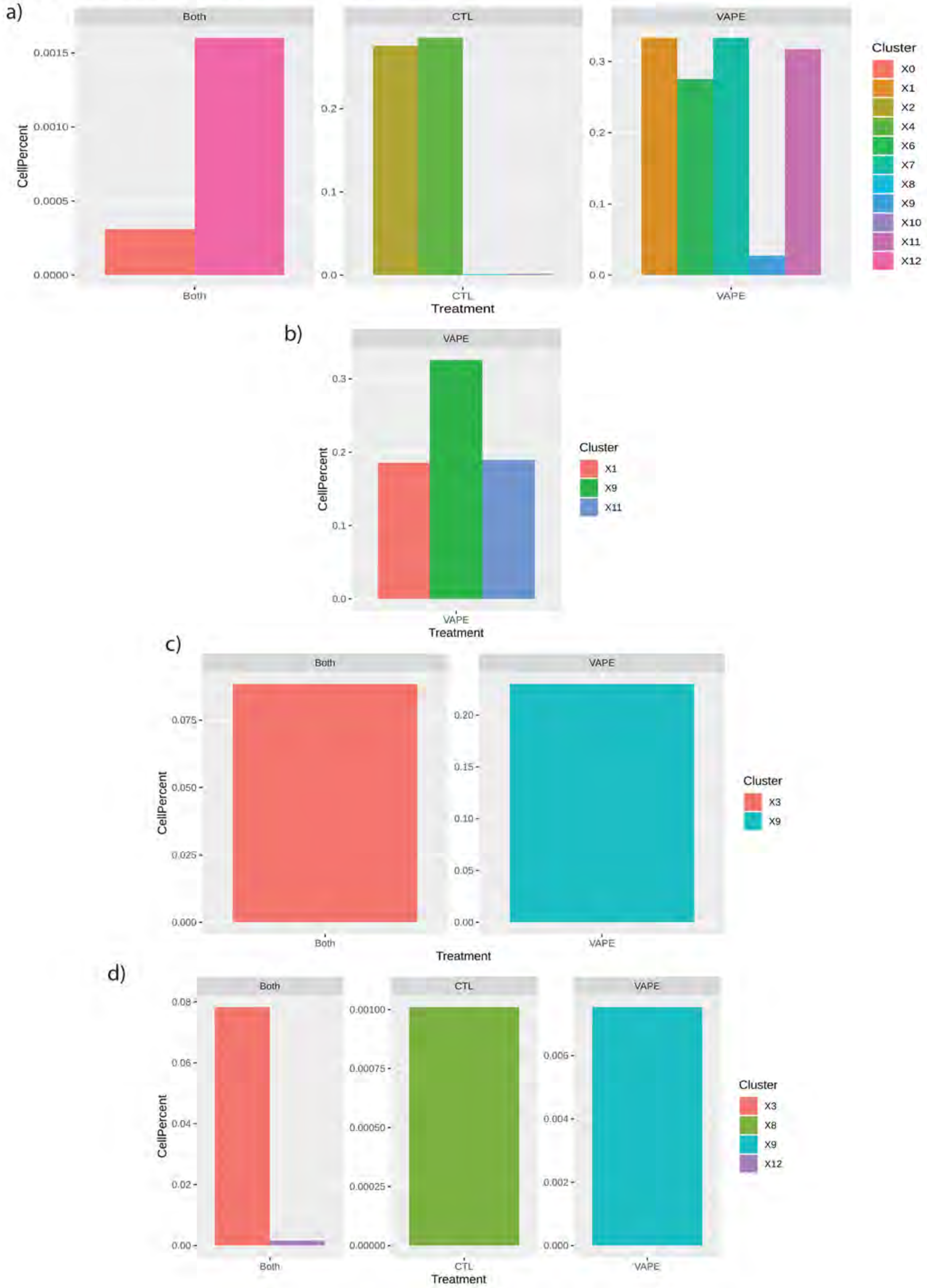

Supplemental Figure 13. Spot contributions by cluster and treatment on cell types mapped to the MCA. a) AT1, b) AT2, c) Ciliated cells and d) Club cells.

### Supplemental Figure 14

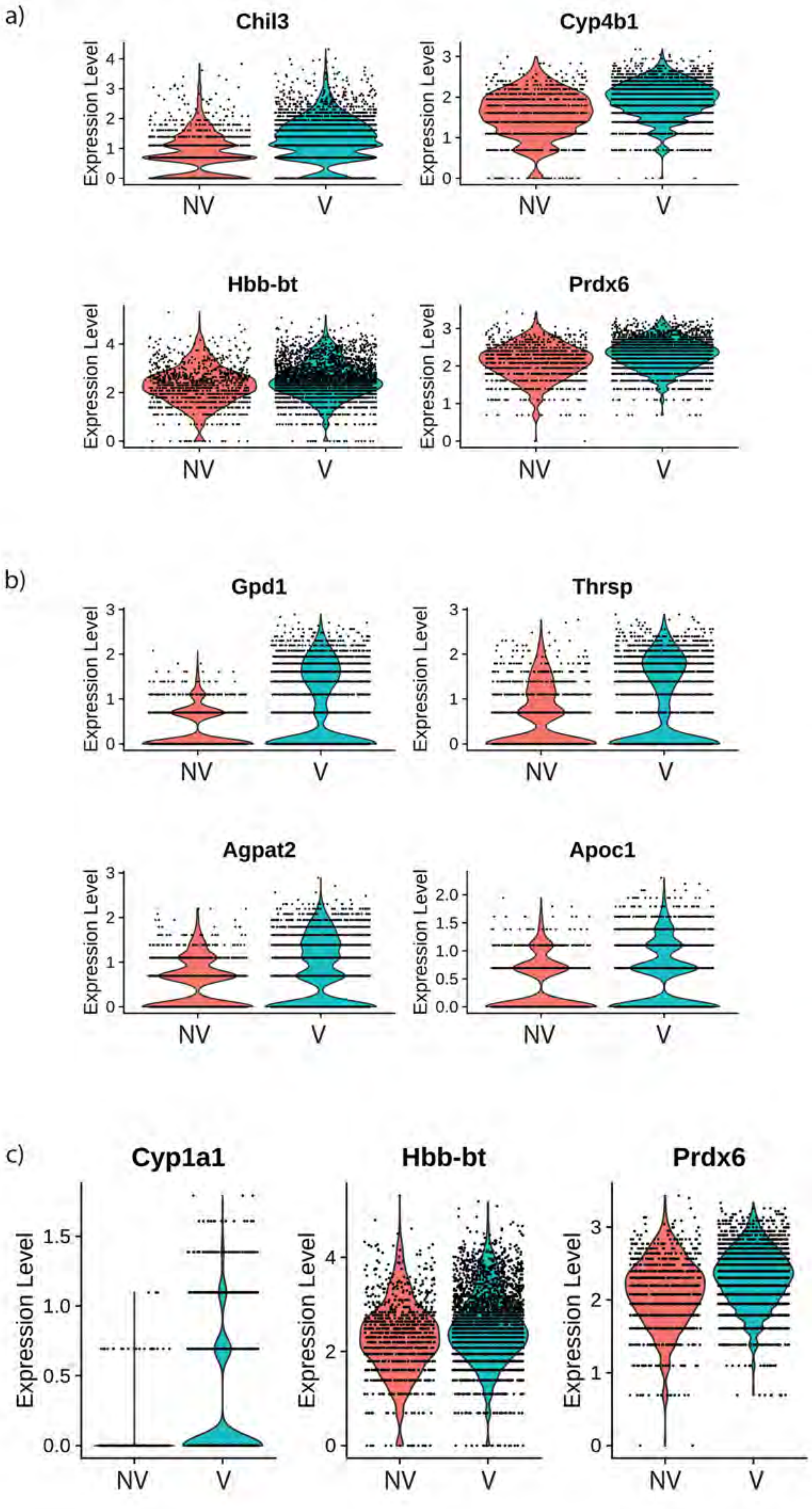

Supplemental Figure 14. Gene targets upregulated on parenchyma of vaped samples grouped by ontology. Violin plots indicating the single spot distribution and expression of gene targets of GO terms: a) drug catabolic process, b) glycerolipid metabolic process and c) hydrogen peroxide metabolic process.

### Supplemental Figure 15

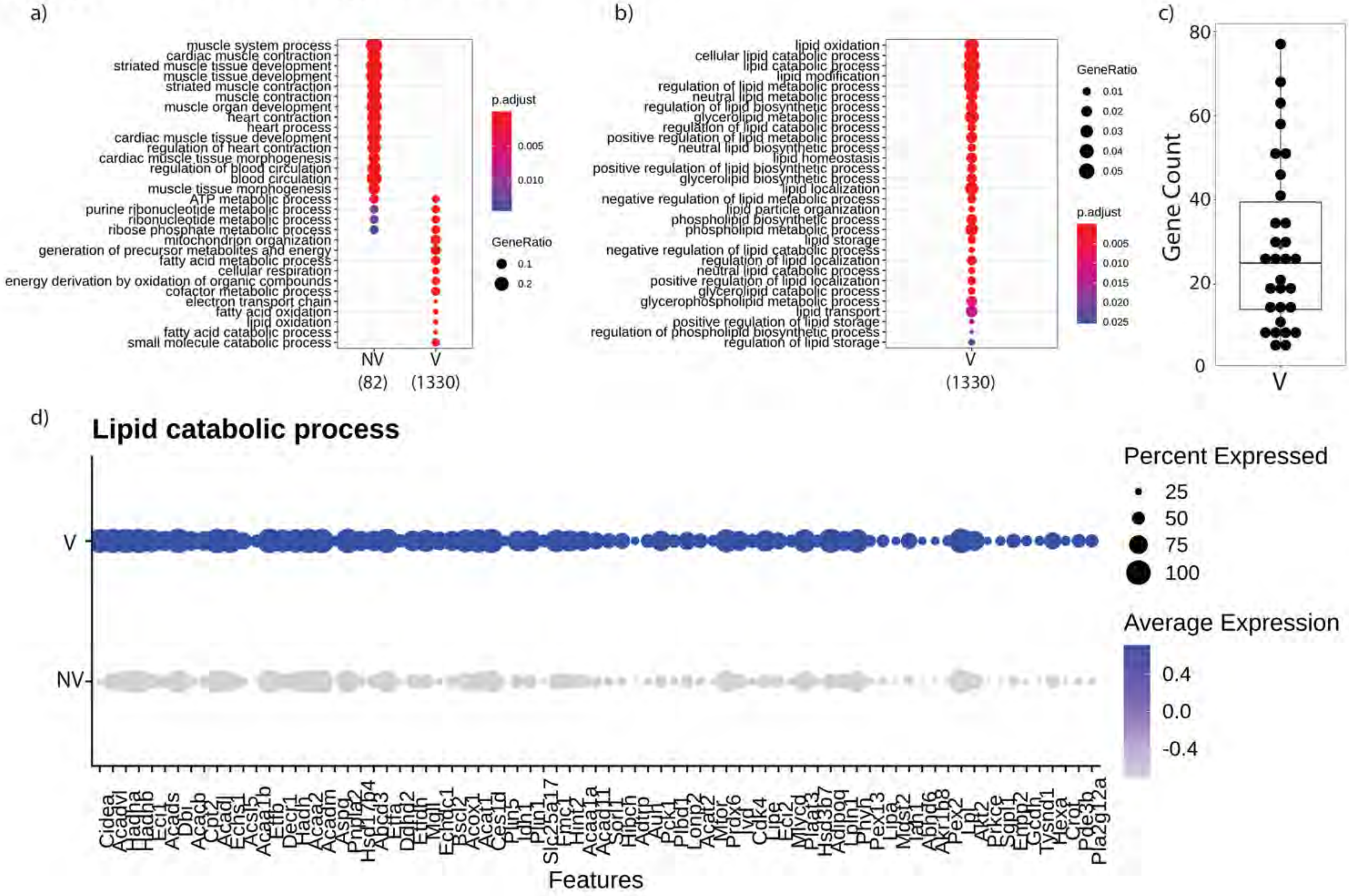

Supplemental Figure 15. Gene targets upregulated on upper airway of vaped samples grouped by ontology. GO terms results from Gene Ontology analysis annotated by Biological Process and presented as a) global analysis and grouped by b) lipid metabolism ontologies on Non-Vaped and Vaped samples. Circle diameter represents the gene ratio, while significance level is color-coded according to heatmap scale. c) Gene counts per GO term grouped by lipid metabolism ontologies on Non-Vaped and Vaped samples. d) Dotplot representing expression of marker of lipid catabolic process GO terms. Circle diameter represents the percentage of spots expressing a particular gene, while normalized average expression is represented by color intensity.

### Supplemental Figure 16

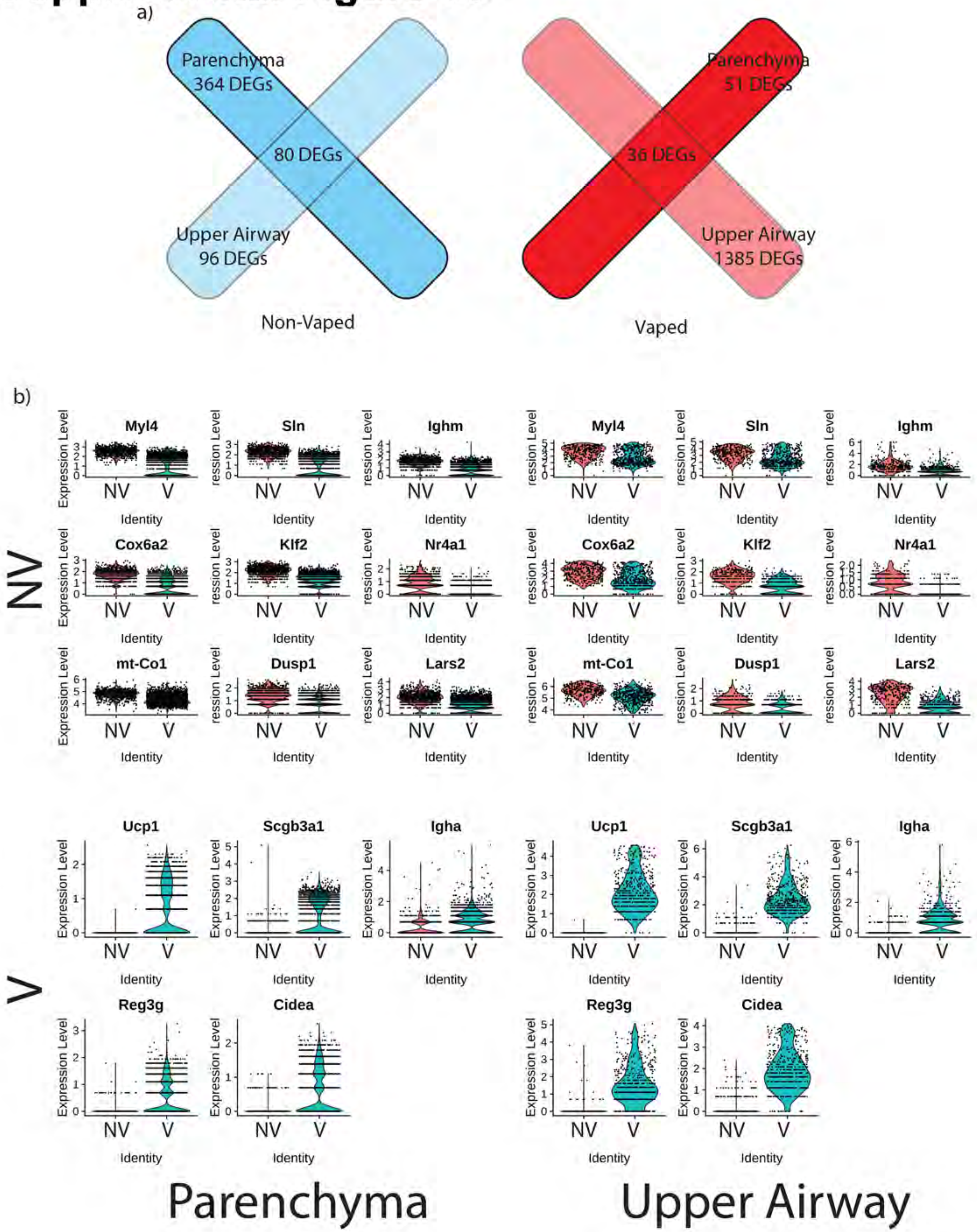

Supplemental Figure 16. Upregulated gene targets intersect on parenchyma and upper airway of non-vaped and vaped samples. a) Venn diagram showing intersection between DEGs of both treatments and pulmonary regions. b) Violin plots indicating the single spot distribution and expression of sample gene targets within DEG intersect.
